# Exploring Various Polygenic Risk Scores for Skin Cancer in the Phenomes of the Michigan Genomics Initiative and the UK Biobank with a Visual Catalog: *PRSWeb*

**DOI:** 10.1101/384909

**Authors:** Lars G. Fritsche, Lauren J. Beesley, Peter VandeHaar, Robert B. Peng, Maxwell Salvatore, Matthew Zawistowski, Sarah A. Gagliano, Sayantan Das, Jonathon LeFaive, Erin O. Kaleba, Thomas T. Klumpner, Stephanie E. Moser, Victoria M. Blanc, Chad M. Brummett, Sachin Kheterpal, Gonçalo R. Abecasis, Stephen B. Gruber, Bhramar Mukherjee

## Abstract

Polygenic risk scores (PRS) are designed to serve as a single summary measure, condensing information from a large number of genetic variants associated with a disease. They have been used for stratification and prediction of disease risk. The construction of a PRS often depends on the purpose of the study, the available data/summary estimates, and the underlying genetic architecture of a disease. In this paper, we consider several choices for constructing a PRS using summary data obtained from various publicly-available sources including the UK Biobank and evaluate their abilities to predict outcomes derived from electronic health records (EHR). We examine the three most common skin cancer subtypes in the USA: basal cell carcinoma, cutaneous squamous cell carcinoma, and melanoma. The genetic risk profiles of subtypes may consist of both shared and unique elements and we construct PRS to understand the common versus distinct etiology. This study is conducted using data from 30,702 unrelated, genotyped patients of recent European descent from the Michigan Genomics Initiative (MGI), a longitudinal biorepository effort within Michigan Medicine. Using these PRS for various skin cancer subtypes, we conduct a phenome-wide association study (PheWAS) within the MGI data to evaluate their association with secondary traits. PheWAS results are then replicated using population-based UK Biobank data. We develop an accompanying visual catalog called *PRSweb* that provides detailed PheWAS results and allows users to directly compare different PRS construction methods. The results of this study can provide guidance regarding PRS construction in future PRS-PheWAS studies using EHR data involving disease subtypes.

**Author summary:** In the study of genetically complex diseases, polygenic risk scores synthesize information from multiple genetic risk factors to provide insight into a patient’s risk of developing a disease based on his/her genetic profile. These risk scores can be explored in conjunction with health and disease information available in the electronic medical records. They may be associated with diseases that may be related to or precursors of the underlying disease of interest. Limited work is available guiding risk score construction when the goal is to identify associations across the medical phenome. In this paper, we compare different polygenic risk score construction methods in terms of their relationships with the medical phenome. We further propose methods for using these risk scores to decouple the shared and unique genetic profiles of related diseases and to explore related diseases’ shared and unique secondary associations. Leveraging and harnessing the rich data resources of the Michigan Genomics Initiative, a biorepository effort at Michigan Medicine, and the larger population-based UK Biobank study, we investigated the performance of genetic risk profiling methods for the three most common types of skin cancer: melanoma, basal cell carcinoma and squamous cell carcinoma.

## Introduction

The underlying risk factors of genetically complex diseases are numerous. Genome-wide association studies (GWAS) on thousands of diseases and traits have made great strides to uncover a vast array of genetic variants that contribute to genetic predispositions to a disease [1]. In order to harness the information from a large number of genetic variants, a popular approach is to summarize their contribution through polygenic risk scores (PRS). While the performance of PRS to predict disease outcomes at a population level has been modest for many diseases including most cancers, PRS have successfully been applied for risk stratification of cohorts [2, 3] and recently have been used to screen a multitude of clinical phenotypes (collectively called the medical phenome) for secondary trait associations [4, 5]. The goal of these phenome-wide screenings is to uncover phenotypes that share genetic components with the primary trait that, if pre-symptomatic, could shed biological insights into the disease pathway and inform early interventions or screening efforts for individuals at risk. However, limited prior work is available guiding the choice of PRS construction for testing associations across the medical phenome.

In the post-GWAS era and with the availability of large biobank data from multiple sources, general guidance for constructing a PRS for a phenotype of interest is needed. A PRS of the general form 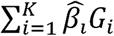 requires specification of three things: a list of markers *G*_l_, *G*_2_, *G*_K_, the depth of the list or the number of markers (*K*), and the choice of the weights 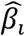 These choices can be based on information extracted from the latest GWAS or GWAS meta-analysis (when available), the NHGRI-EBI GWAS catalog of published results [1] (when available), or summary data for GWAS corresponding to each phenotype, e.g., from efforts that comprehensively screened the UK Biobank (UKB) phenome [6, 7]. While various methods of constructing PRS have been widely studied for predicting the primary phenotype collected through population-based sampling [8, 9], it is unknown how the different PRS will be associated with a multitude of other diagnoses across the medical phenome. This study attempts to bridge this knowledge gap.

In this paper, we first explore strategies for constructing a PRS using markers and weights obtained from either the latest GWAS or the NHGRI-EBI GWAS catalog that have reached genome-wide significance. We compare the PRS in terms of their performance [10] for the three most common skin cancer subtypes in the USA: basal cell carcinoma (MIM: 614740) [11], cutaneous squamous cell carcinoma [12] and melanoma (MIM: 155601) [13]. We compare the two strategies using an independent biobank of genetic, demographic, and phenotype data collected by the Michigan Genomics Initiative (MGI), a longitudinal biorepository effort within Michigan Medicine (University of Michigan) [4, 14]. Based on these results, we choose a PRS construction strategy for each skin cancer subtype for further analysis.

For the chosen PRS corresponding to each skin cancer subtype, we perform a phenome-wide association study (PheWAS) relating the PRS to the electronic health record (EHR)-based phenome of MGI. We call such a study a PRS-PheWAS.^4^ PRS-PheWAS results are then replicated using the population-based UK Biobank data. In order to identify secondary associations that are not driven by the primary phenotype, we perform an additional “exclusion” PRS-PheWAS for each skin cancer subtype in which we exclude subjects with any type of observed skin cancer.^4^ These studies demonstrate differences in PheWAS results for PRS constructed for particular disease subtypes and the ability of such studies to reproduce known associations between secondary phenotypes and particular disease subtypes.

We then describe an approach for using PRS to (1) understand the shared and unique genetic architecture of disease subtypes and to (2) identify shared and unique secondary phenotype associations related to this genetic architecture. We define a new PRS for each skin cancer subtype using loci ***unique*** to that subtype’s chosen PRS. We further construct a composite PRS for general skin cancer consisting of loci ***common*** among all subtypes’ PRS. While merging distinct clinical entities into a compound PRS may seem counterintuitive in terms of specificity, such an approach may increase power to identify dermatological features through PheWAS that are shared by all three subtypes, which may in turn provide guidance for general skin cancer screening efforts and sun protection behavior.

The NHGRI-EBI GWAS Catalog and Latest GWAS PRS construction methods are based on published GWAS studies, which only report risk variants that reached genome-wide significance (usually defined by a P-value threshold of P < 5×10^−8^). However, it is likely that there are additional risk variants below this threshold that could be associated with the trait but have not reached statistical significance [15]. Incorporating non-significant variants may conceivably improve the predictive power of a PRS but may also add additional random false positive signals, which in turn could dilute the discriminatory power of the true risk variants and diminish any predictive gain [8, 16]. To explore whether a PRS constructed using additional non-significant loci may outperform a PRS using only loci reaching genome-wide significance, we evaluated a PRS constructed using publicly available genome-wide summary statistics from the UK Biobank at six different p-value thresholds both in terms of associations with skin cancer phenotypes and in terms of secondary phenotype associations. There is an extensive literature on constructing genome-wide PRS using random effects, shrinkage methods, or thresholding (our focus) [17-19], but none of these methods have been evaluated in a PheWAS setting.

In this paper, we focus our attention on skin cancer, but the approaches used in this paper can be applied to study many other phenotypes. We chose to use skin cancer as a demonstrative example for a variety of reasons. First, our discovery dataset (MGI) is particularly enriched for skin cancer cases due to the strong skin cancer clinical program at Michigan Medicine and due to the high rate of surgery for skin cancer patients. MGI primarily recruits participants undergoing surgery and is therefore enriched for cancers and other medical comorbidities when compared to a general population [4]. Additionally, skin cancer has well-defined subtypes, which allows us to explore subtype-specific PRS constructed for several related but distinct diseases in terms of their performance for related skin cancer outcomes. Skin cancer also provides a setting in which there may be genetic factors uniquely related to particular subtypes as well as genetic factors that are shared risk factors for all skin cancer subtypes. The various PRS construction methods explored in this paper delivered tools to explore shared and subtype-specific phenotypes and may provide an enhanced understanding of the genome x phenome landscape.

We develop an online visual web catalog called *PRSweb* that provides PRS-PheWAS results for melanoma, basal cell carcinoma, and squamous cell carcinoma. PheWAS results are available using three different PRS construction methods explored in this paper: Latest GWAS, NHGRI-EBI GWAS Catalog, and the UK Biobank GWAS summary statistics using different significance thresholds. The weights and the marker list for each PRS method can be downloaded. Furthermore, PheWAS summary statistics can be accessed from *PRSweb* (see **Web resources**), providing future investigators with readily available and useful tools to perform further analyses.

Comprehensive phenome-wide and genome-wide analyses of large biobank studies with publicly available summary statistics can be rich resources for PRS construction, especially if the trait-of-interest’s prevalence is high in the biobank. Using PRS, we can synthesize complex genetic information that is then used to identify these shared genetic components across phenotypes. Compared to prior and existing literature, our contribution is new in four principal directions: (1) comparing various PRS construction methods in terms of their relationships with related EHR-derived phenotypes (2) comparing PRS associations with secondary phenotypes across the phenome of MGI (academic medical center) and UK Biobank (population-based), (3) developing PRS-based methods for understanding the shared and unique genetic contribution across disease sub-types both in terms of disease biology and in terms of secondary phenotype associations, and (4) introducing a publicly accessible online visual catalog to visually represent the genome x phenome landscape and access summary data from GWAS and PheWAS.

## Material and methods

### Discovery and replication cohorts

#### MGI cohort (discovery cohort)

Participants were recruited through the Michigan Medicine health system while awaiting diagnostic or interventional procedures either during a preoperative visit prior to the procedure or on the day of the procedure that required anaesthesia. Opt-in written informed consent was obtained. In addition to coded biosamples and secure protected health information, participants understood that all EHR, claims, and national data sources – linkable to the participant – may be incorporated into the MGI databank. Each participant donated a blood sample for genetic analysis, underwent baseline vital signs and a comprehensive history and physical assessment. Data were collected according to Declaration of Helsinki principles. Study participants’ consent forms and protocols were reviewed and approved by local ethics committees (IRB ID HUM00099605). In the current study, we report results obtained from 30,702 unrelated, genotyped samples of recent European ancestry with available integrated EHR data (∼90 % of all MGI participants were inferred to be of recent European ancestry) [4].

#### UK Biobank cohort (replication cohort)

The UK Biobank is a population-based cohort collected from multiple sites across the United Kingdom and includes over 500,000 participants aged between 40 and 69 years when recruited in 2006–2010 [20]. The open access UK Biobank data used in this study included genotypes, ICD9 and ICD10 codes, inferred sex, inferred white British-European ancestry, kinship estimates down to third degree, birthyear, genotype array, and precomputed principal components of the genotypes.

### Genotyping, sample quality control and imputation

#### MGI

DNA from 37,412 blood samples was genotyped on customized Illumina Infinium CoreExome-24 bead arrays and subjected to various quality control filters that resulted in a set of 392,323 polymorphic variants. Principal components and ancestry were estimated by projecting all genotyped samples into the space of the principal components of the Human Genome Diversity Project reference panel using PLINK (938 unrelated individuals) [21, 22]. Pairwise kinship was assessed with the software KING [23], and the software fastindep was used to reduce the data to a maximal subset that contained no pairs of individuals with 3rd-or closer degree relationship [24]. We also removed patients not of recent European descent from the analysis, resulting in a final sample of 30,702 unrelated subjects. Additional genotypes were obtained using the Haplotype Reference Consortium using the Michigan Imputation Server [25] and included over 17 million imputed variants with R^2^<0.3 and/or minor allele frequency (MAF) <0.1%. Genotyping, quality control and imputation are described in detail elsewhere [4]. **Table 1** provides some descriptive statistics of the MGI and UK Biobank samples.

**Table 1.**
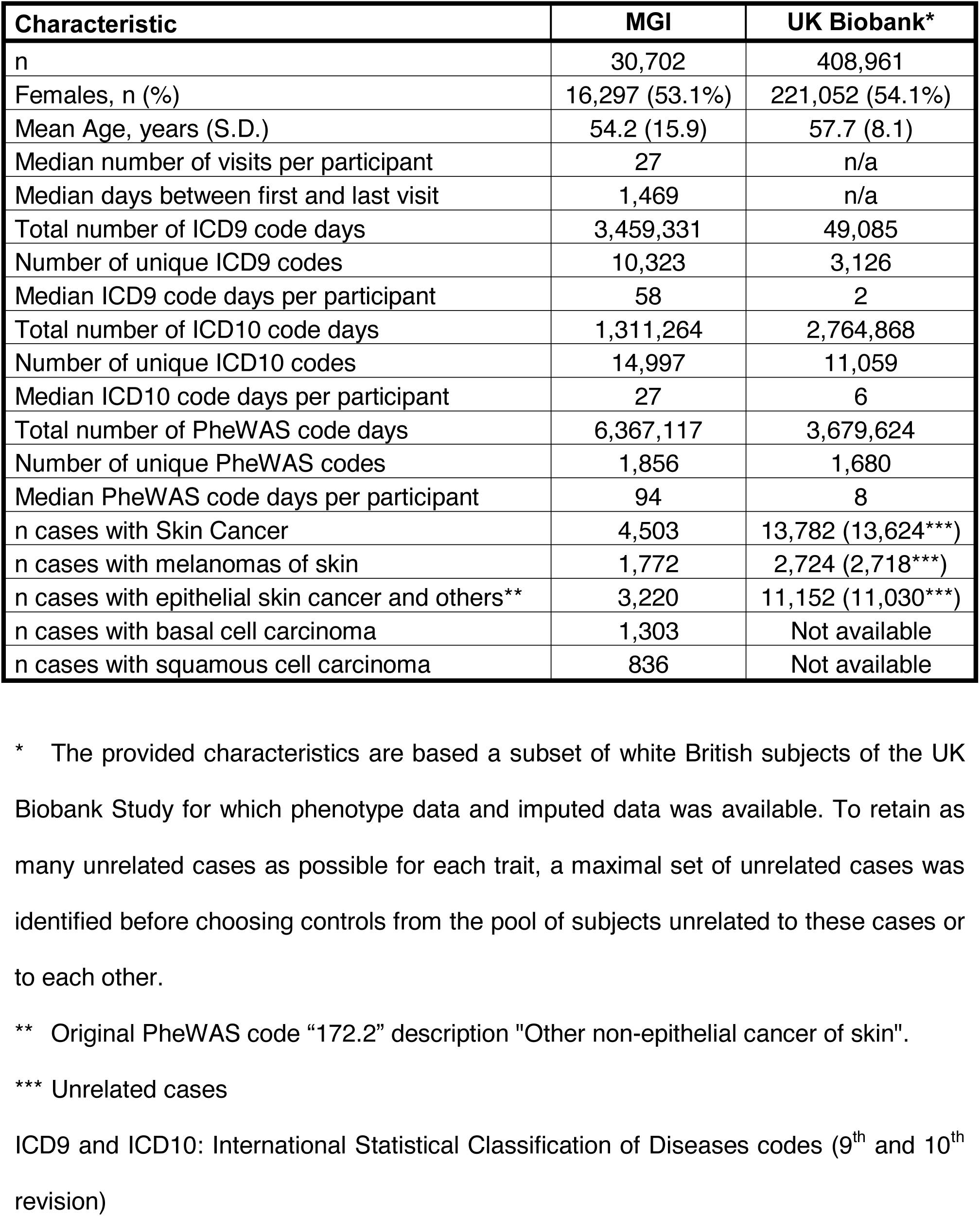
Demographics and Clinical Characteristics of the Analytic Datasets

| Characteristic | MGI | UK Biobank* |
| --- | --- | --- |
| n | 30,702 | 408,961 |
| Females, n (%) | 16,297 (53.1%) | 221,052 (54.1%) |
| Mean Age, years (S.D.) | 54.2 (15.9) | 57.7 (8.1) |
| Median number of visits per participant | 27 | n/a |
| Median days between first and last visit | 1,469 | n/a |
| Total number of ICD9 code days | 3,459,331 | 49,085 |
| Number of unique ICD9 codes | 10,323 | 3,126 |
| Median ICD9 code days per participant | 58 | 2 |
| Total number of ICD10 code days | 1,311,264 | 2,764,868 |
| Number of unique ICD10 codes | 14,997 | 11,059 |
| Median ICD10 code days per participant | 27 | 6 |
| Total number of PheWAS code days | 6,367,117 | 3,679,624 |
| Number of unique PheWAS codes | 1,856 | 1,680 |
| Median PheWAS code days per participant | 94 | 8 |
| n cases with Skin Cancer | 4,503 | 13,782 (13,624***) |
| n cases with melanomas of skin | 1,772 | 2,724 (2,718***) |
| n cases with epithelial skin cancer and others** | 3,220 | 11,152 (11,030***) |
| n cases with basal cell carcinoma | 1,303 | Not available |
| n cases with squamous cell carcinoma | 836 | Not available |

\* The provided characteristics are based a subset of white British subjects of the UK Biobank Study for which phenotype data and imputed data was available. To retain as many unrelated cases as possible for each trait, a maximal set of unrelated cases was identified before choosing controls from the pool of subjects unrelated to these cases or to each other.
\*\* Original PheWAS code “172.2” description "Other non-epithelial cancer of skin".
\*\*\* Unrelated cases
ICD9 and ICD10: International Statistical Classification of Diseases codes (9<sup>th</sup> and 10<sup>th</sup> revision)

#### UK Biobank

The UK Biobank is a population-based cohort collected from multiple sites across the United Kingdom [20]. After quality control, we phased and imputed the 487,409 UK Biobank genotyped samples against the Trans-Omics for Precision Medicine (TOPMed) reference panel (see **Web resources**), which is composed of 60,039 multi-ethnic samples and 239,756,147 SNP and indel variants sequenced at high depth (30x). The phasing step was carried out on 81 chromosomal chunks with around 10,000 genotyped variants in each chunk using the software Eagle (with the “kbpwt” parameter set at 80,000) [26]. The imputation was carried out in 137 chromosomal chunks of around 20 Mbp in length with Mbp of total overlap on either side using the imputation tool Minimac4 (see **Web resources**). To increase computational efficiency, we imputed each of the chunks in batches of 10,000 samples at a time and then merged them back using BCFtools. Since Minimac4 imputes each sample independently, analyzing our samples in batches did not change their imputation estimates. However, this sampling would result in different imputation quality estimates for each batch, and thus we collapsed the estimates to generate imputation quality estimates across all the study samples. After imputation, we filtered out variants with estimated imputation accuracy of R^2^ < 0.1, which left us with 177,895,992 variants.

### Phenome generation

#### MGI

The MGI phenome was used as the discovery dataset and was based on the Ninth and Tenth Revision of the International Statistical Classification of Diseases (ICD9 and ICD10) code data for 30,702 unrelated, genotyped individuals of recent European ancestry. These ICD9 and ICD10 codes were aggregated to form up to 1,857 PheWAS traits using the PheWAS R package (as described in detail elsewhere[4, 27]). For each trait, we identified case and control samples. To minimize differences in age and sex distributions or extreme case-control ratios as well as to reduce computational burden, we matched up to 10 controls to each case using the R package “MatchIt” [28]. Nearest neighbor matching was applied for age and PC1-4 (using Mahalanobis-metric matching; matching window caliper/width of 0.25 standard deviations) and exact matching was applied for sex and genotyping array. A total of 1,578 case control studies with >50 cases were used for our analyses of the MGI phenome.

#### UK Biobank

The UK Biobank phenome was used as a replication dataset and was based on ICD9 and ICD10 code data of 408,961 white British [14], genotyped individuals that were aggregated to PheWAS traits in a similar fashion (as described elsewhere [7]). To remove related individuals and to retain larger sample sizes, we first selected a maximal set of unrelated cases for each phenotype (defined as no pairwise relationship of 3^rd^ degree or closer [24, 29]) before selecting a maximal set of unrelated controls unrelated to these cases. Similar to MGI, we matched up to 10 controls to each case using the R package “MatchIt” [28]. Nearest neighbor matching was applied for birthyear and PC1-4 (using Mahalanobis-metric matching; matching window caliper/width of 0.25 standard deviations) and exact matching was applied for sex and genotyping array. 1,366 case control studies with >50 cases each were used for our analyses of the UK Biobank phenome.

Additional phenotype information for MGI and UK Biobank is included in **S1 Text Fig B** and **S2 Text Tables F-H**.

### Risk SNP selection

For each skin cancer subtype (melanoma, basal cell carcinoma, and squamous cell carcinoma), we generated three different sets of PRS: (1) based on merged summary statistics published in the NHGRI EBI GWAS catalog [1], (2) based on the latest available GWAS meta-analysis [30-32] and (3) based on publicly available GWAS summary statistics from the UK Biobank data [7].

#### GWAS Catalog SNP selection

We downloaded previously reported GWAS variants from the NHGRI-EBI GWAS Catalog (file date: February 28, 2018) [1, 33]. None of the currently available skin cancer discovery studies included in the catalog used any subset of the MGI cohort or data from the UK Biobank. Single nucleotide polymorphism (SNP) positions were converted to GRCh37 using variant IDs from dbSNP: build 150 (UCSC Genome Browser) after updating outdated dbSNP IDs to their merged dbSNP IDs. Entries with missing risk alleles, risk allele frequencies, or odds ratios were excluded. If a reported risk allele did not match any of the reported forward strand alleles of a non-ambiguous SNP (not A/T or C/G) in the imputed genotype data (which correspond to the alleles of the imputation reference panel), we assumed minus strand designation and corrected the effect allele to its complementary base of the forward strand. Entries with a reported risk allele that did not match any of the alleles of an ambiguous SNP (A/T and C/G) in our data were excluded at this step. We only included entries with broad European ancestry (as reported by the NHGRI-EBI GWAS Catalog). As a quality control check, we compared the reported risk allele frequencies (RAF) in controls with the RAF of 14,770 MGI individuals who had no cancer diagnosis (for chromosome X variants, we calculated RAF in females only). We then excluded entries whose RAF deviated more than 15%. This chosen threshold is subjective and was based on clear differentiation between correct and likely flipped alleles on the two diagonals (see **S1 Text Fig A**) as noted frequently in GWAS meta-analyses quality control procedures [34]. For each analyzed cancer type, we extracted risk variants that were also present in our genotype data and estimated pairwise linkage disequilibrium (LD; correlation r^2^) using the allele dosages of the corresponding controls. For pairwise correlated SNPs (r^2^>0.1) or SNPs with multiple entries, we kept the SNP with the most recent publication date (and smaller *P* value, if necessary) and excluded the other (**S2 File Table I**).

#### Selection of risk SNPs from largest GWAS

In a similar fashion, we extracted and filtered reported association signals from large GWAS meta-analyses on basal cell carcinoma [31], cutaneous squamous cell carcinoma [30] and melanoma [32] (**S2 File Table I**).

#### Genome-wide SNP selection of UK-Biobank-based GWAS

We obtained GWAS summary statistics for the ICD9- and ICD10-based PheWAS codes “172” (skin cancer; 13,752 cases versus 395,071 controls), “172.11” (melanoma; 2,691 cases versus 395,071 controls), and “172.2” (non-epithelial skin cancer; 11,149 cases versus 395,071 controls) from a public download [7] (see **Web resources**). These GWAS analyzed up to 408,961 white British European-ancestry samples with generalized mixed model association tests that used the saddlepoint approximation to calibrate the distribution of score test statistics and thus could control for unbalanced case-control ratios and sample relatedness [7]. For each trait, we reduced these summary statistics to SNPs that were reported with minor allele frequencies > 0.5% and were also available for the MGI data. Next, we performed linkage LD clumping of all variants with p-values < 5×10^−4^ using the imputed allele dosages to obtain independent risk SNPs (LD threshold of r^2^ > 0.1 and a maximal SNP distance of 1 Mb). We limited the LD calculations to 10,000 randomly selected, unrelated, white British individuals to reduce the computational burden. Finally, we created subsets of these independent SNPs with p-values <5×10^−9^, <5×10^−8^, <5 × 10^−7^, <5×10^−6^, <5×10^−5^, and <5×10^−4^ (**S2 File Table J**).

### Construction of the polygenic risk scores

For each of the obtained SNP sets for each trait, we constructed a PRS as the sum of the allele dosages of risk increasing alleles of the SNPs weighted by their reported log odds ratios. Restated, the PRS for subject j in MGI was of the form PRS_*j*_=∑_i_ (β_*i*_ G_*ij*_ where *i* indexes the included loci for that trait, (β_i_ is the log odds ratios retrieved from the external GWAS summary statistics for locus *i,* and G_*ij*_ is a continuous version of the measured dosage data for the risk allele on locus *i* in subject *j*. The PRS variable was created for each MGI and UKB participant. For comparability of effect sizes corresponding to the continuous PRS across cancer traits and PRS construction methods, we transformed each PRS to the standard Normal distribution using “ztransform” of the R package “GenABEL” [35].

### Statistical analysis

In this study, we constructed PRS for three skin cancer subtypes using two different PRS construction methods (using the Latest GWAS or the corresponding entries of the GWAS Catalog). To compare the association between PRS and skin cancer phenotypes across different PRS construction methods, we fit the following model for each PRS and skin cancer phenotype:

logit (P(Phenotype is present | PRS, Age, Sex, Array, PC)) =(β_O_ + (β_PRS_ PRS + (β_Age_Age + (β_Sex_ Sex + (β_Array_Array + β PC, where the PCs were the first four principal components obtained from the principal component analysis of the genotyped GWAS markers and where “Array” represents the genotyping array. Our primary interest is in (β_PRS_, while the other factors (Age, Sex and PC) were included to address potential residual confounding and do not provide interpretable estimates due to the preceding application of case control matching. Firth’s bias reduction method was used to resolve the problem of separation in logistic regression (Logistf in R package “EHR”) [36-38], a common problem for binary or categorical outcome models when for a certain part of the covariate space there is only one observed value of the outcome, which often leads to very large parameter estimates and standard errors.

We then evaluated each PRS’s (1) ability to discriminate between cases and controls by determining the area under the receiver-operator characteristics (ROC) curve (AUC) using R package “pROC” [39]; (2) calibration using Hosmer-Lemeshow Goodness Of Fit test of the R package “ResourceSelection” [40, 41]; and (3) accuracy with the Brier Score of R package “DescTools” [42]. These evaluations did not adjust for additional covariates. We used these metrics and the logistic regression results to choose a PRS construction method to use for each skin cancer subtype moving forward. To explore the impact of incorporating non-significant loci into the PRS construction, we further performed the above analyses with PRS constructed using UK Biobank GWAS summary statistics with different p-value thresholds.

Using the chosen PRS for each subtype, we conducted two PheWAS to identify other phenotypes associated with the PRS first for the 1,578 phenotypes in MGI and then for the 1,366 phenotypes from UK Biobank. To evaluate PRS-phenotype associations, we conducted Firth bias-corrected logistic regression by fitting a model of the above form for each phenotype and data source. Age represents the birth year in UK Biobank. To adjust for multiple testing, we applied the conservative phenome-wide Bonferroni correction according to the analyzed PheWAS codes (n_MGI_ = 1,578 or n_UK Biobank_ = 1,366). In Manhattan plots, we present –log10 (*p*-value) corresponding to tests of H_O_: (β_PRS_ = 0. Directional triangles on the PheWAS plot indicate whether a phenome-wide significant trait was positively (pointing up) or negatively (pointing down) associated with the PRS.

To investigate the possibility of the secondary trait associations with PRS being completely driven by the primary trait association, we performed a second set of PheWAS after excluding individuals affected with the primary or related cancer traits for which the PRS was constructed, referred to as “exclusion PRS PheWAS” as described previously [4]. We then constructed new PRS scores representing shared and subsite-unique genetic components and performed a PheWAS for each.

To evaluate how well prior presence of an identified secondary non-skin-cancer diagnosis can identify subjects with increased risk of developing skin cancer, we created a binary variable taking the value 1 if a given subject (1) was diagnosed with the non-skin-cancer diagnosis and then diagnosed with skin cancer at least 365 days after or (2) was diagnosed with the non-skin-cancer diagnosis and never diagnosed with skin cancer. We then fit a Firth bias-corrected logistic regression of the following form: logit (P(Primary phenotype is present | Predictor, Age, Sex, Array, PC))=(β_0_ + (β_PRS_ *I* (Secondary non skin cancer trait) + (β_Age_ Age + (β_Sex_ Sex + (β_Array_ Array+ β PC where Array and PC were defined as before. Unless otherwise stated, analyses were performed using R 3.4.4 [43].

### Development of an online visual catalog: *PRSweb*

The online open access online visual catalog *PRSweb* available at https://statgen.github.io/PRSweb was implemented using “Pandas”, a Data Analysis Library, which offers high level of performance for large data structures and data analysis in the Python3 environment [44]. In combination with “Jinja2”, a templating language for Python, and “Bootstrap”, a Cascading Style Sheets (CSS) framework (see Web resources), static HTML files were compiled and allow easy and fast hosting of all PRS-PheWAS results. The interactive plots are drawn with the JavaScript library “LocusZoom.js” (see Web resources) offers dynamic plotting, automatic plot sizing and label positioning.

## Results

### Assessing various PRS construction methods

We first explored the comparative performance of two PRS construction strategies in terms of the resulting PRS associations with related phenotypes in the skin cancer setting. **Table 2** provides the results.

**Table 2.** Associations of constructed PRS with skin cancer traits in MGI

| PRS<br>(Number of<br>SNPs) |  | Skin Cancer<br>n = 4,503 | Melanoma<br>n = 1,896 | Basal Cell<br>Carcinoma<br>n = 1,303 | Squamous Cell<br>Carcinoma<br>n = 836 |
| --- | --- | --- | --- | --- | --- |
| *PRS based on GWAS Catalog* |  |  |  |  |  |
| Melanoma<br>(29) | PRS OR<sup>a</sup><br>P-value<sup>a</sup><br>AUC<sup>b</sup><br>HL $\chi^2$, P-value<sup>c</sup><br>Brier Score | 1.41 (1.35,1.47)<br>2.7x10<sup>-53</sup><br>0.57 (0.56,0.58)<br>10,0.24<br>0.14 | **1.68 (1.57,1.79)**<br>1.3x10<sup>-53</sup><br>**0.61 (0.60,0.62)**<br>5.3,0.72<br>0.09 | 1.42 (1.31,1.53)<br>7.3x10<sup>-19</sup><br>0.57 (0.56,0.59)<br>12,0.16<br>0.091 | 1.3 (1.19,1.44)<br>4.3x10<sup>-08</sup><br>0.55 (0.53,0.57)<br>3.7,0.89<br>0.09 |
| Basal cell<br>carcinoma<br>(32) | PRS OR<sup>a</sup><br>P-value<sup>a</sup><br>AUC<sup>b</sup><br>HL $\chi^2$, P-value<sup>c</sup><br>Brier Score | 1.39 (1.33,1.44)<br>8x10<sup>-60</sup><br>0.57 (0.56,0.58)<br>13,0.12<br>0.14 | 1.37 (1.29,1.45)<br>4.8x10<sup>-25</sup><br>0.57 (0.56,0.58)<br>8.6,0.38<br>0.091 | **1.82 (1.70,1.95)**<br>3.6x10<sup>-65</sup><br>**0.64 (0.62,0.65)**<br>9.5,0.3<br>0.09 | 1.4 (1.28,1.52)<br>1.4x10<sup>-14</sup><br>0.57 (0.55,0.59)<br>13,0.11<br>0.09 |
| Squamous cell<br>carcinoma<br>(18) | PRS OR<sup>a</sup><br>P-value<sup>a</sup><br>AUC<sup>b</sup><br>HL $\chi^2$, P-value<sup>c</sup><br>Brier Score | 1.28 (1.24,1.33)<br>4.8x10<sup>-42</sup><br>0.56 (0.56,0.57)<br>4.7,0.79<br>0.14 | 1.35 (1.28,1.43)<br>2x10<sup>-28</sup><br>0.58 (0.56,0.59)<br>5.3,0.72<br>0.091 | **1.39 (1.31,1.48)**<br>7.9x10<sup>-26</sup><br>**0.59 (0.57,0.60)**<br>5.1,0.75<br>0.091 | 1.29 (1.19,1.39)<br>1.8x10<sup>-10</sup><br>0.56 (0.54,0.59)<br>7.8,0.46<br>0.09 |
| *PRS based on Latest GWAS* |  |  |  |  |  |
| Melanoma<br>(20) | PRS OR<sup>a</sup><br>P-value<sup>a</sup><br>AUC<sup>b</sup><br>HL $\chi^2$, P-value<sup>c</sup><br>Brier Score | 1.48 (1.41,1.55)<br>3.5x10<sup>-55</sup><br>0.57 (0.56,0.58)<br>3,0.93<br>0.14 | **1.78 (1.65,1.92)**<br>7x10<sup>-53</sup><br>**0.61 (0.59,0.62)**<br>6.7,0.56<br>0.09 | 1.60 (1.47,1.75)<br>7.9x10<sup>-27</sup><br>0.59 (0.57,0.60)<br>2.5,0.96<br>0.091 | 1.38 (1.24,1.53)<br>4x10<sup>-09</sup><br>0.56 (0.54,0.58)<br>4.3,0.83<br>0.09 |
| Basal cell<br>carcinoma<br>(28) | PRS OR<sup>a</sup><br>P-value<sup>a</sup><br>AUC<sup>b</sup><br>HL $\chi^2$, P-value<sup>c</sup><br>Brier Score | 1.42 (1.36,1.48)<br>5.8x10<sup>-61</sup><br>0.58 (0.57,0.58)<br>4.3,0.83<br>0.14 | 1.43 (1.34,1.52)<br>7x10<sup>-29</sup><br>0.58 (0.56,0.59)<br>16,0.051<br>0.091 | **1.84 (1.71,1.97)**<br>2.8x10<sup>-60</sup><br>**0.63 (0.62,0.65)**<br>4,0.86<br>0.09 | 1.45 (1.32,1.58)<br>1.2x10<sup>-15</sup><br>0.57 (0.55,0.60)<br>17,0.035<br>0.09 |
| Squamous cell<br>carcinoma<br>(10) | PRS OR<sup>a</sup><br>P-value<sup>a</sup><br>AUC<sup>b</sup><br>HL $\chi^2$, P-value<sup>c</sup><br>Brier Score | 1.44 (1.38,1.5)<br>1.1x10<sup>-70</sup><br>0.58 (0.57,0.59)<br>17,0.027<br>0.14 | 1.54 (1.45,1.64)<br>2.9x10<sup>-46</sup><br>0.60 (0.58,0.61)<br>13,0.13<br>0.09 | **1.62 (1.52,1.73)**<br>1.8x10<sup>-43</sup><br>**0.61 (0.60,0.63)**<br>6,0.64<br>0.09 | 1.52 (1.39,1.65)<br>2.1x10<sup>-21</sup><br>0.59 (0.57,0.61)<br>4.9,0.76<br>0.09 |

<sup>a</sup> Association of each cancer with continuous PRS that were transformed to standard normal distribution. Point estimates, 95% confidence intervals and P- values are obtained by fitting Firth's Bias-Corrected Logistic Regression.
<sup>b</sup> Area under the curve of the receiver operating characteristic (ROC) curve with 95% confidence intervals.
<sup>c</sup> Hosmer-Lemeshow Goodness-of-Fit Test

#### Comparisons within methods

Using the GWAS Catalog construction method, the melanoma PRS was more strongly associated with and had better discrimination for the melanoma phenotype than the other skin cancer phenotypes. For the PRS based on the GWAS Catalog, the odds ratio (OR) of the melanoma PRS was 1.68 (95% CI, [1.57, 1.79]). By “discrimination,” we refer to the ability of the PRS to distinguish melanoma cases and controls, which is measured by AUC. The melanoma PRS AUC for the melanoma phenotype is 0.61 (95 % CI, [0.60, 0.62]). Similarly, the basal cell carcinoma PRS was most strongly associated with and had the best discrimination for the basal cell carcinoma phenotype, with an OR of 1.82 (95% CI, [1.70, 1.95]) and an AUC of 0.64 (95% CI, [0.62, 0.65]). Unlike the other cancer subtypes, the squamous cell carcinoma PRS did not appear to be most strongly associated with the squamous cell carcinoma phenotype. Instead, it was most strongly associated with and most discriminative for basal cell carcinoma. For all three skin cancer subtypes, the PRS produced higher Brier scores for overall skin cancer, suggesting that the subtype-defined PRS were less accurate for predicting skin cancer as a whole. We obtain similar conclusions for the Latest GWAS method.

#### Comparisons across methods

For each cancer subtype, we compared the PRS-subtype associations for the two PRS construction methods. **Melanoma**: For the melanoma PRS, the GWAS Catalog method and the Latest GWAS method produced similar performance in terms of AUC, OR, Hosmer-Lemeshow goodness of fit, and Brier score. For example, the AUC for melanoma for the GWAS Catalog melanoma PRS was 0.61 (95% CI, [0.60, 0.62]). The corresponding AUC for the Latest GWAS method was 0.61 (95% CI, [0.59, 0.62]). **S1 Text Fig J** compares PRS weights to corresponding SNP-melanoma associations in MGI and UK Biobank. **Basal Cell Carcinoma**: As with melanoma, the basal cell carcinoma PRS produced similar results under the GWAS Catalog and Latest GWAS construction methods. The basal cell carcinoma AUC under the GWAS catalog method was 0.64 (95% CI, [0.62, 0.65]) and the AUC under the Latest GWAS method was 0.63 (95% CI, [0.62, 0.65]). The OR values and Brier score values were nearly identical, and neither approach produced evidence of lack of fit based on the Hosmer-Lemeshow statistic. **Squamous Cell Carcinoma:** The squamous cell carcinoma PRS was not more strongly associated with the squamous cell carcinoma phenotype than the other phenotypes. However, we do observe that the squamous cell carcinoma phenotype using the GWAS Catalog method (0.56, 95% CI [0.54, 0.59]) produced a lower AUC compared to the Latest GWAS method (0.59, 95% CI [0.57, 0.61]). While a difference of 0.03 may not seem like a large difference in AUC in other applications, any improvement in AUC for PRS associations with observed phenotypes may be considered appreciable [45]. These two methods produced identical Brier scores, and the Latest GWAS method resulted in a stronger association between the PRS and the squamous cell carcinoma phenotype (OR of 1.29, 95% CI [1.19, 1.39] vs OR of 1.52, 95% CI [1.39, 1.65]).

Using the above comparisons between the two PRS construction methods, we chose a single PRS construction method for each skin cancer subtype to use in subsequent analyses. For melanoma and basal cell carcinoma, we chose the GWAS Catalog method. While the GWAS Catalog and Latest GWAS methods were very similar for these two subtypes, we chose to pursue the GWAS Catalog PRS for future analysis due to the larger number of loci for these PRS (29 vs 20 for melanoma and 32 vs 28 for basal cell carcinoma). We choose the Latest GWAS method for squamous cell carcinoma due to its improved AUC over the GWAS Catalog method. We will denote the chosen PRS for melanoma, basal cell carcinoma, and squamous cell carcinoma as mPRS, bPRS, and sPRS respectively.

### PheWAS using the chosen PRS in MGI

Using each of the chosen PRS described above (mPRS, bPRS, and sPRS), we tested the association between each PRS and each of the 1,578 constructed phenotypes in MGI. For each PRS, the strongest associations were observed with dermatologic neoplasms that included overall skin cancer, melanoma, “other non-epithelial cancer of skin” (the PheWAS over-category of basal and squamous cell carcinoma), and carcinoma in situ of skin. In addition, secondary dermatologic traits such as actinic keratosis (with over-category “degenerative skin conditions and other dermatoses”), chronic dermatitis due to solar radiation (with over-category “dermatitis due to solar radiation”), and seborrheic keratosis were found to be associated with all three PRS (**Fig 1** and **S2 File Table K**). mPRS was most strongly associated with the melanoma phenotype (OR 1.67, 95% CI [1.56, 1.79]), while bPRS was most strongly associated with carcinoma in situ of the skin (OR 1.51, 95% CI [1.39, 1.64]) followed closely by “non-epithelial cancer of the skin” (OR 1.47, 95% CI [1.41, 1.54]). sPRS was most strongly associated with carcinoma in situ of the skin (OR 1.79, 95% CI [1.65, 1.94]). The OR of all these phenotypes indicated an increased risk for primary and secondary traits with increasing PRS.

**Fig 1.**
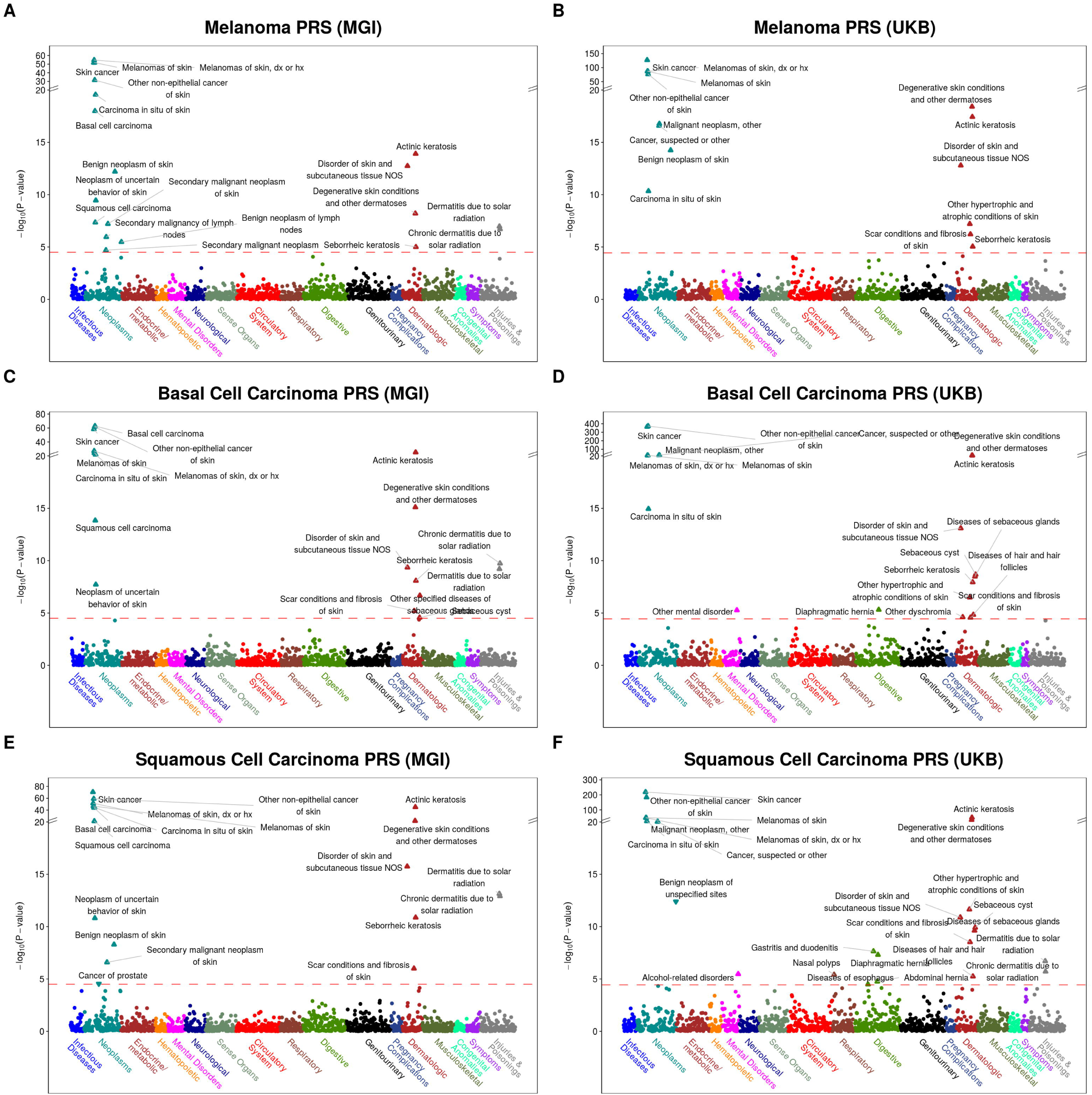
PRS-PheWAS in MGI and UKB phenomes. The horizontal line indicates phenome-wide significance.

### Validation of PRS-PheWAS in UK Biobank

To substantiate the detected dermatologic associations, we reiterated the association screen of the three PRS in the matched phenome of the population-based UK Biobank data set (**Fig 1**). In general, stronger evidence for association was found in UKB compared to MGI. This may be driven by the larger sample sizes, e.g. a total of 13,623 skin cancer cases versus 4,503 in MGI. In the UK Biobank phenome, the large majority of the previous associations with dermatologic neoplasms were validated with the exception of the trait “dermatitis due to solar radiation”, which had substantially fewer cases in UKB compared to MGI (390 versus 2,959 cases). Unlike MGI, all three PRS were significantly associated (at the phenome-wide level) with “cancer, suspected or other” and “malignant neoplasm, other.”

### Exclusion PheWAS using the chosen PRS in MGI

In order to explore whether the identified PRS-phenotype associations were driven by the primary trait used to define the PRS (for example, as a side effect of treatment given after diagnosis with the primary trait), we performed a PheWAS for each PRS in which we excluded subjects who were cases for the primary trait or other skin cancer subtypes [4]. Results are shown in **S2 File Table K** and **S1 Text Fig C**. Actinic keratosis, a skin condition believed to be a precursor to non-melanoma skin cancers, remained significantly associated with the squamous cell carcinoma PRS in MGI and all three PRS in UK Biobank [46][47, 48]. No other phenotypes were significant for MGI. “Sebaceous cyst” and its over-category “diseases of the sebaceous gland” were significant in the main UK Biobank PheWAS and remained significantly associated with basal cell carcinoma PRS and squamous cell carcinoma PRS in UK Biobank in the Exclusion PheWAS.

### Sub-analysis of actinic keratosis as a predictor of future skin cancer

Actinic keratosis (AK) is a rough, scaly patch of skin that usually develops after years of cumulative skin exposure [49]. Previous research has identified actinic keratosis as a common pre-malignant condition for squamous cell carcinoma (SCC) [46]. Actinic keratosis has also been identified as a potential precursor to basal cell carcinoma (BCC) [47, 48]. The availability of temporal information of diagnoses in the MGI cohort offered the opportunity to explore actinic keratosis as a potential precursor for development of skin cancer in MGI.

**Fig 2** shows the ROC curves and AUC values for diagnosis of actinic keratosis at least one year before any skin cancer diagnosis and its association with future BCC or SCC diagnosis. AK diagnosis alone has little discrimination abilities, with AUC values of 0.52 (95% CI [0.51, 0.53]) for BCC and 0.51 (95% CI [0.50, 0.61]) for SCC. The bPRS and sPRS provide comparatively good discrimination SCC (AUC 0.63 [0.62, 0.65] for BCC and 0.59 [0.57, 0.61] for SCC). The combination of prior AK diagnosis and bPRS provided further improvement in discrimination, with an AUC of 0.65 (95% CI [0.64, 0.67]).

**Fig 2.**
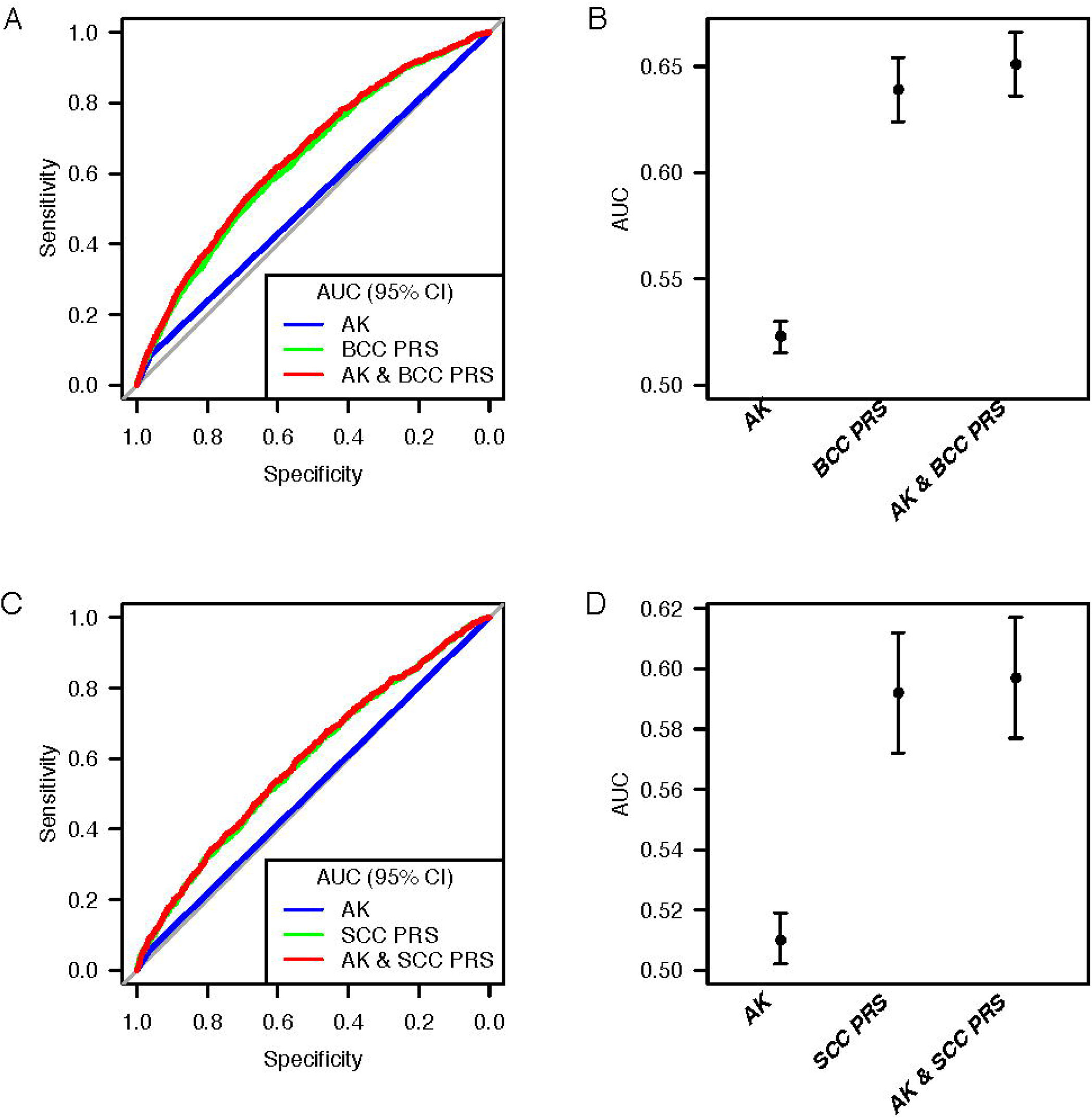
Comparison of predictors. Actinic keratosis (AK), at least 365 prior to any skin cancer diagnosis as predictor for basal cell carcinoma (BCC) (A and B) and squamous cell carcinoma (SCC) (C and D). The PRS for BCC and SCC as well as the combined predictors are shown for comparison.

**S1 Text Tables A and B** provides odds ratio estimates relating AK and the PRS to future BCC and SCC diagnosis. In unadjusted models, the odds of BCC diagnosis were significantly higher in subjects with a prior actinic keratosis diagnosis (OR 1.46, 95% CI [1.18, 1.80]). Notably, when we adjust for both bPRS and AK diagnosis, the unadjusted and adjusted effects of both variables are similar, suggesting that AK diagnosis may be an independent predictor of future BCC diagnosis. In contrast, AK diagnosis was **not** an independent predictor of SCC diagnosis. **S1 File Fig G** shows the timing of an AK diagnosis relative to a skin cancer diagnosis for patients with both diagnoses. For subjects with basal cell carcinoma or squamous cell carcinoma, AK diagnoses tended to occur prior to the skin cancer diagnosis (often within 8 years).

### PRS-PheWAS for shared and unique loci across skin cancer subtypes

In the PRS-PheWAS analyses, we note a striking overlap in the secondary dermatological traits significantly associated with each of the three PRS (mPRS, bPRS, sPRS). One potential explanation for this is that subjects may have more screening after an initial skin cancer diagnosis. Indeed, many subjects have multiple skin cancer diagnoses (**S1 Text Fig D**). **Fig 3** shows the number of risk loci shared by different PRS. Six risk loci are shared between the mPRS, bPRS, and sPRS.

**Fig 3.**
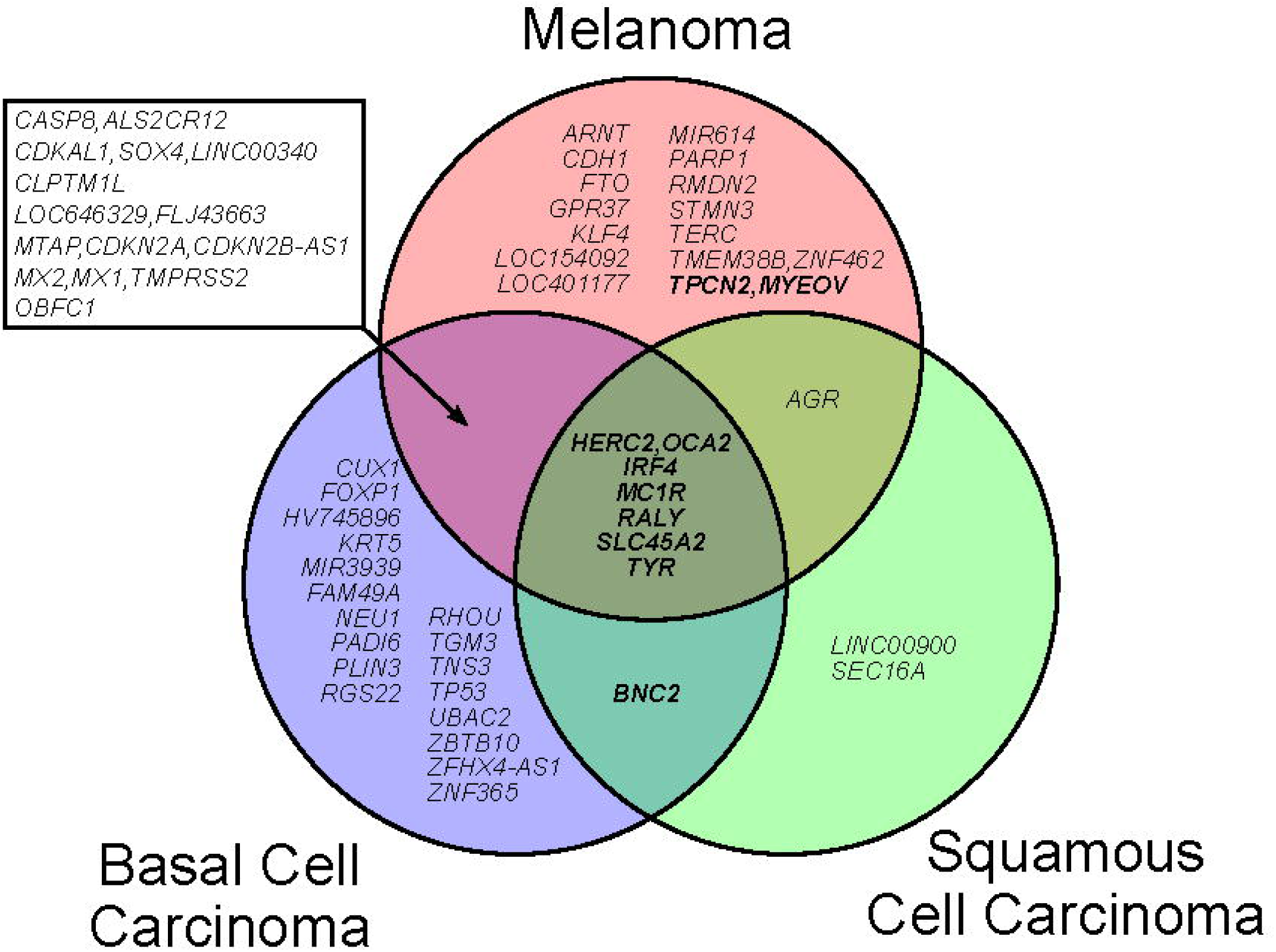
Overlap between the three skin cancer trait loci. Reported risk SNPs within 1 Mb were merged into the same locus. Loci that were also reported to be associated with skin tanning ability are highlighted in bold. Loci were named according to the closest RefSeq genes (except *M1CR* a 385 kb locus with 16 RefSeq genes and *HV745896* named after a nearby, uncurated mRNA sequence).

**Fig 4.**
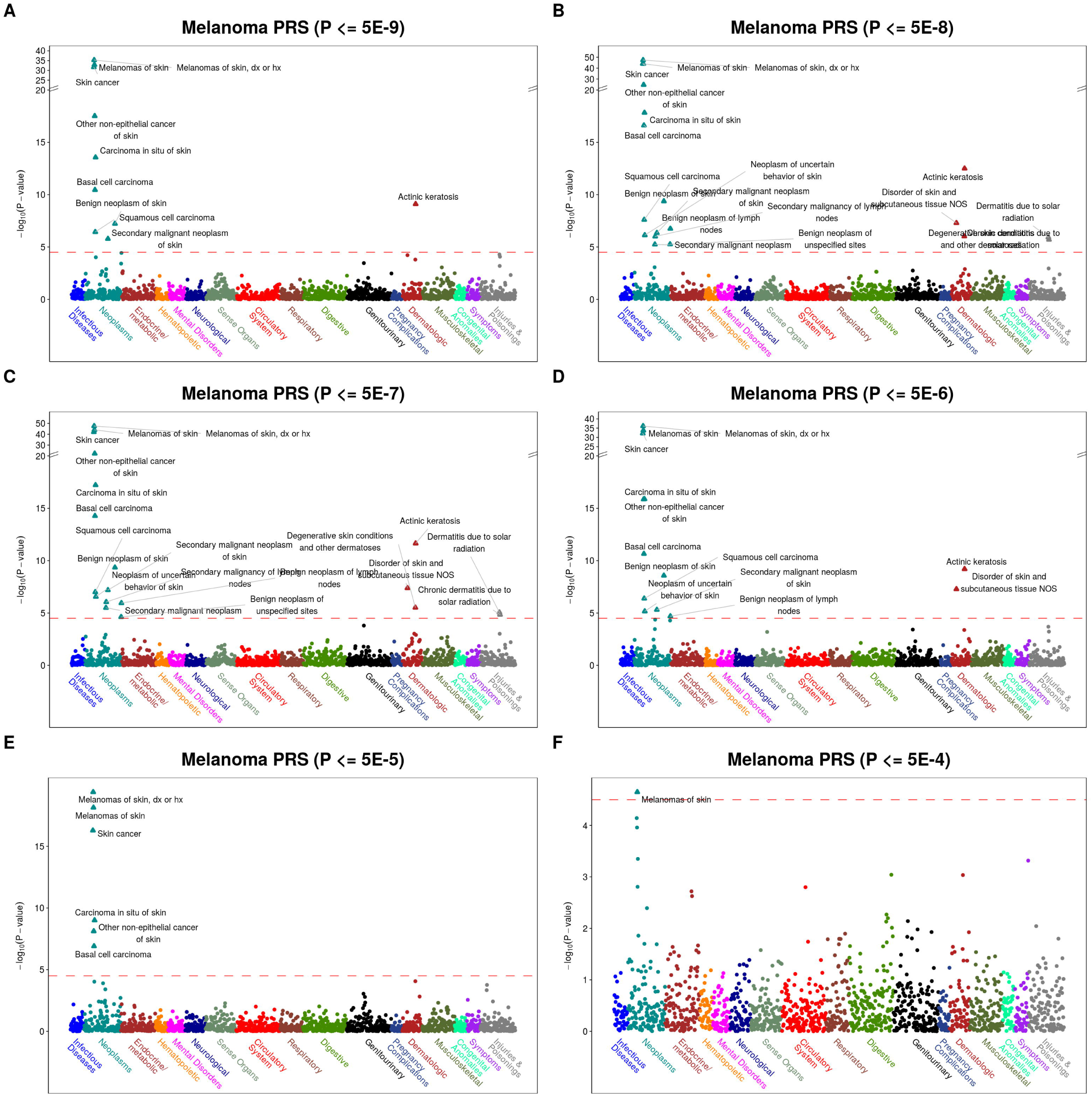
PheWAS on melanoma PRS constructed using UK Biobank statistics at different depths. Results are shown with increasing depth from (A – F): P <= 5×10^−9^, 5×10^−8^, 5×10^−7^, 5×10^−6^, 5×10^−5^, 5×10^−4^.

This observation inspired follow-up exploration in which we defined a PRS for each cancer subtype using the loci unique to that subtype’s chosen PRS. We call these new PRS scores mPRS-u, bPRS-u, and sPRS-u, which reflect the unique loci in the PRS for melanoma, basal cell carcinoma, and squamous cell carcinoma respectively. We also define a PRS consisting of all loci shared across the three skin cancer subtypes, which we call the shared PRS.

**S1 Text Table C** shows the association between the various constructed PRS and the skin cancer phenotypes. As with mPRS, mPRS-u was most strongly associated with the melanoma phenotype and is not significantly associated with the other skin cancer subtypes. The bPRS-u score was similarly most strongly associated with basal cell carcinoma and not significantly associated with the other subtypes. We note that the melanoma AUC for the mPRS score was 0.61 (95% CI, [0.60, 0.62]) and is only 0.55 (95% CI, [0.53, 0.55]) for the mPRS-u score. Similarly, the basal cell carcinoma AUC for the bPRS score was 0.64 (95% CI, [0.62, 0.65]) and is only 0.57 (95% CI, [0.55, 0.58]) for the bPRS-u score. The sPRS-u score is not more strongly associated with the squamous cell carcinoma phenotype than the other skin cancer subtypes. For this reason, we do not include this PRS in further analyses. The shared PRS constructed as the unweighted sum of risk alleles of loci present in all three PRS scores (mPRS, bPRS, and sPRS) is more strongly associated with all three subtype phenotypes than the overall skin cancer phenotype, and the overall skin cancer phenotype also has the lowest AUC and highest Brier score.

**S1 Text Fig E** shows PRS-PheWAS results using mPRS-u and bPRS-u. The scores again reveal their subtype specificity in both phenomes, while no secondary associations were observed. Although not shown here, additional exploration into the loci identified uniquely for each subtype may provide some insight into subtype-specific biological mechanisms. **S1 Text Fig F** shows PRS-PheWAS results for the shared PRS. Most strikingly, the shared skin cancer PRS was associated with the top skin cancer and dermatologic traits that were previously found to be associated with the three partially overlapping PRS constructs, suggesting that a shared genetic risk may be driving many of these secondary associations. These six underlying loci (*HERC2* [MIM 605837] /*OCA2* [MIM 611409], *IRF4* [MIM 601900], *MC1R* [MIM 155555], *RALY* [MIM 614663], *SLC45A2* [MIM 606202] and *TYR* [MIM 606933]) were previously found to be associated not only with skin cancer traits, but also with pigmentation traits of skin, eyes and hair (**Fig 3;** MIM 266300) [31, 50-69].

One of these pigmentation traits, skin tanning ability, the tendency of skin to sunburn rather than to suntan, is a well-known risk factor for all skin cancer traits [69, 70]. A PRS based on the independent risk variants of a recent GWAS meta-analysis on skin tanning ability [70] was strongly associated with overall skin cancer, melanoma, basal cell carcinoma, and squamous cell carcinoma and even outperformed the constructed PRS of the former two traits (**S1 Text Table C**). Furthermore, the skin tanning ability PRS PheWAS identified a very similar set of traits as the shared skin cancer PRS but in general revealed stronger associations (**S1 Text Fig F**).

### PRS construction based on UK Biobank summary statistics

The NHGRI-EBI GWAS Catalog and Latest GWAS PRS construction methods are based on published GWAS studies, which often only report risk variants that reached genome-wide significance, but we may believe that incorporating additional risk variance below this threshold may improve predictive power of a PRS. To explore whether a PRS incorporating non-significant loci will outperform a PRS incorporating only significant loci, we constructed PRS using loci related to the phenotype at six different p-value thresholds based on publicly available GWAS summary statistics from the UK Biobank. Larger p-values indicate greater SNP depth (with more SNPs being incorporated into the PRS).

The collection of UK Biobank GWAS results did not include basal cell carcinoma or squamous cell carcinoma subtypes; rather, it included only the merged trait ‘non-epithelial cancer of skin’ (**S1 Text Fig B**). Thus, we limited our assessment of the summary statistics to the overall skin cancer GWAS (UKB PheWAS code “172”: 13,752 skin cancer cases versus 395,071 controls) and the melanoma GWAS (UKB PheWAS code “172.11”: 2,691 melanoma cases versus 395,071 controls) (**S2 File Table J**).

**S1 Text Table D** provides the results. As with the other PRS construction methods, the melanoma PRS was most strongly associated with and discriminative for the melanoma phenotype for all p-value cutoffs except 5×10^−4^. For this p-value cutoff, the melanoma PRS had similar AUC and OR for the melanoma and basal cell carcinoma phenotypes. This p-value cutoff represents the least conservative inclusion cutoff with 1,193 included loci, and its results indicated that inclusion of too many suggestive SNPs at lower thresholds may reduce PRS performance. However, we also note that the most conservative cutoff (5×10^−9^) produced a PRS with only six loci and a weaker OR and AUC compare to other PRS created with less stringent cutoffs. Like the other PRS construction methods, the melanoma PRS was less accurate for predicting overall skin cancer compared to the individual skin cancer subtypes. The best performance in terms of AUC and OR relating to the melanoma phenotype were observed for p-value thresholds 5×10^−7^ and 5×10^−8^, which included 13 and 9 loci respectively. The small number of loci identified by this method at more conservative p-value cutoffs may be driven by the lower sample size for melanoma in the UK Biobank compared to the published melanoma GWAS meta-analyses (n cases = 2,691 and n cases = 6,628, respectively). We note that the melanoma PRS constructed using the UK Biobank summary statistics produced lower AUC across all p-value thresholds than was seen for the Latest GWAS and GWAS Catalog PRS construction methods.

The PRS constructed for overall skin cancer was most strongly associated with and discriminative for basal cell carcinoma across all p-value thresholds, with AUCs ranging from 0.59 (95% CI [0.57, 0.60]) to 0.64 (95% CI [0.62, 0.66]) and odds ratios ranging from 1.42 (95% CI [1.33, 1.51]) to 1.73 (95% CI [1.63, 1.84]). The overall skin cancer PRS had the highest Brier score for overall skin cancer, indicating that the overall skin cancer PRS was more accurate at predicting the skin cancer subtypes compared to overall skin cancer. The overall skin cancer PRS had very similar association with and discrimination abilities for the overall skin cancer phenotype across all p-value thresholds except the least conservative (p = 5×10^−4^), for which the AUC and odds ratio were smaller. Overall, the highest AUCs and strongest OR signals for both PRS and all skin cancer phenotypes were found at depths of 5×10^−7^ and 5×10^−8^.

In addition to associations with the primary phenotype, we explored associations between PRS constructed at various UK Biobank summary statistic depths and secondary phenotypes. **Fig 3** (melanoma) and **S1 Text Fig H** (overall skin cancer) show PRS-PheWAS results in MGI using PRS constructed at different depths. As shown in **S1 Text Table E and Fig I**, depths of 5×10^−7^ and 5×10^−8^ produced very similar results, and other depths identified fewer phenotypes associated with the corresponding PRS. Phenotypes that were associated with the PRS at other depths had weaker associations than those observed at 5×10^−7^ and 5×10^−8^.

### Online visual catalog: *PRSweb*

For comparison of the aforementioned PRS-PheWAS results and to provide researchers with resources for future PRS-based analyses, we developed an open access, online visual catalog *PRSweb* available at https://statgen.github.io/PRSweb that enables interactive exploration of the PheWAS results for each of the skin cancer subtypes under each of three different PRS construction methods explored in this paper, for both the MGI and UK Biobank phenomes. *PRSweb* shows PRS-PheWAS plots with various choices of PRS in the drop-down menu (example screenshot in **Fig 5**) and offers downloadable PRS constructs (list of independent risk variants with corresponding weights). Mouse-over boxes offer detailed information about top results if needed, without impeding the overall user experience (grey box in **Fig 5**). Enrichment of cases in the tail of the PRS distribution are presented in interactive forest plots.

**Fig 5.**
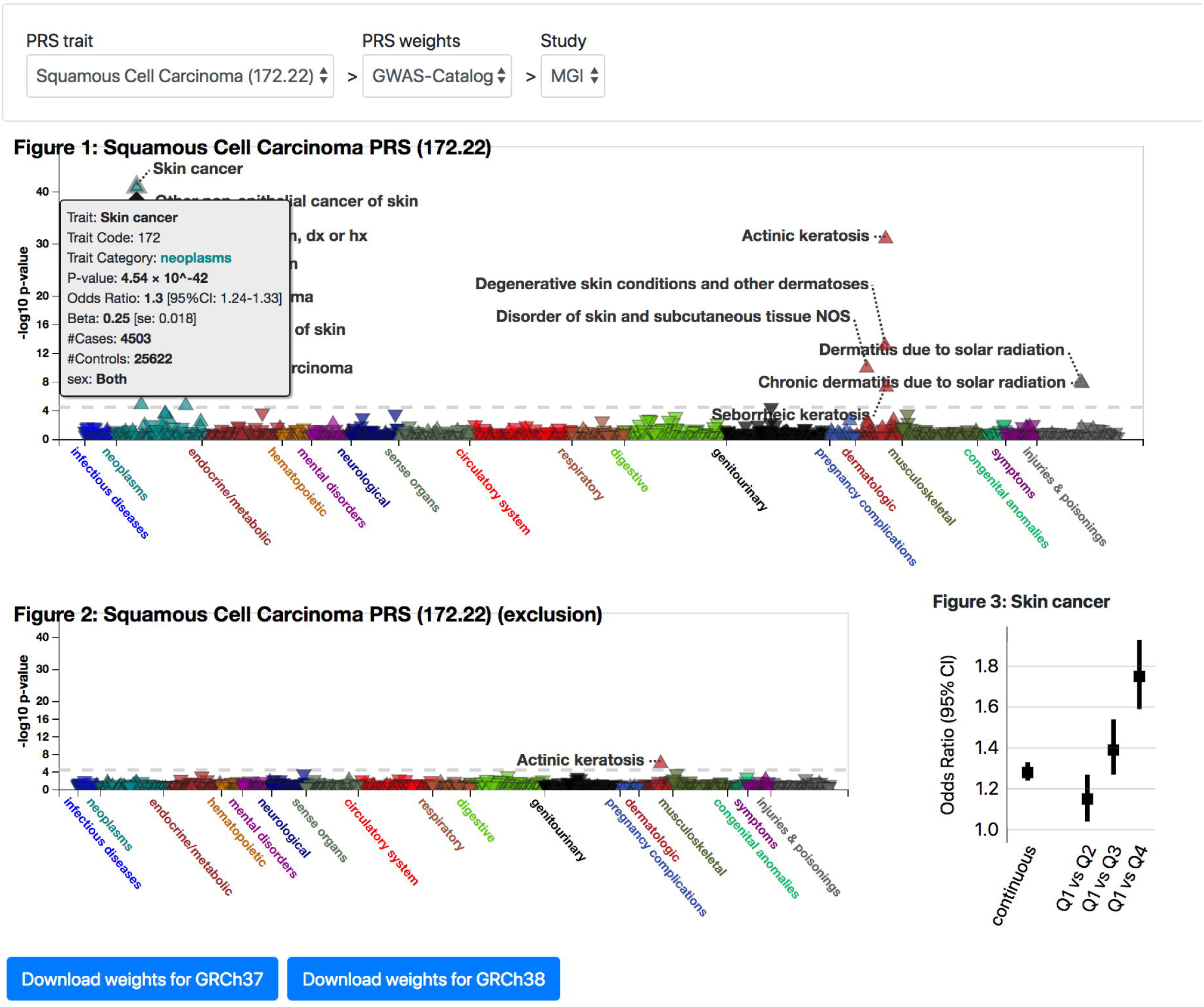
Example view from PRSweb. A selection menu on top allows selection of PRS constructs and phenome while interactive plots with “PheWAS results”, “Exclusion PheWAS results”, and “Associations between PRS and Selected Phenotype” are generated after selection.

## Discussion

PRS combine information from a large number of genetic variants to stratify subjects in terms of their risk for developing a particular disease. However, there are currently no general guidelines for how to construct a PRS for a given *EHR-derived* phenotype. In this paper, we explore strategies for constructing a PRS using markers and weights obtained from various publicly-available sources. First, we consider PRS constructed using markers and weights identified in either (1) the latest GWAS or GWAS meta-analysis or (2) the NHGRI-EBI GWAS Catalog. We compare these two PRS construction methods in terms of their associations with EHR-derived phenotypes for the three most common skin cancer subtypes in the USA: basal cell carcinoma, cutaneous squamous cell carcinoma, and melanoma.

A priori, we may have some belief that the latest (and often the largest) GWAS may provide a better source of evidence to use for PRS construction due to larger sample sizes and (potentially) more carefully curated data. The Latest GWAS and GWAS Catalog methods produced PRS with similar performance in terms of their associations with and discrimination for the primary phenotype used to construct the PRS for both basal cell carcinoma and melanoma. Generally, PRS constructed for melanoma and basal cell carcinoma were most strongly associated with and discriminative for their target phenotypes, indicating that both PRS construction methods were able to provide a higher degree of specificity for the intended skin cancer subtype. In contrast, the PRS for squamous cell carcinoma were not more strongly associated with the squamous cell carcinoma phenotype compared to other skin cancer phenotypes. This may suggest a need for further exploration into genetic factors uniquely related to the squamous cell carcinoma subtype.

For each skin cancer subtype, we performed a PRS-PheWAS to identify secondary phenotypes that are associated with the corresponding PRS. We generally identified many dermatological features in addition to the primary phenotype, indicating the ability of PRS to reproduce associations with the primary phenotype even after multiple testing corrections and covariate adjustment. The majority of these associations were replicated in a PRS-PheWAS performed for the UK Biobank phenome. Our analyses identified actinic keratosis, which is believed to be a precursor to squamous cell and basal cell carcinoma, as an independent predictor of basal cell and squamous cell carcinoma, and we demonstrated that incorporating the PRS in addition to clinical information improved discrimination for future skin cancer diagnoses [46-48].

In an additional analysis, we identified loci that were shared among all three skin cancer subtypes’ PRS. Loci overlap between the PRS for the three subtypes may indicate factors related to common biology between the subtypes. We noted that all shared loci (*HERC2*/*OCA2, IRF4, MC1R, RALY, SLC45A2* and *TYR*) were also loci that had been associated with human pigmentation traits and/or harbor key genes of the biochemical pathway of melanogenesis [50, 54-62, 64, 67-71]. We constructed PRS using SNPs shared by all three skin cancer subtypes and a PRS for skin tanning ability using results from a recent GWAS meta-analysis.[70] The skin tanning ability PRS PheWAS identified a very similar set of traits to the shared PRS PheWAS, suggesting that the shared genetic component may in part represent genetic factors influencing the skin pigmentation and the skin reaction to sun exposure. However, the PRS that are unique to subtypes did not show such a common pathway or mechanism.

The Latest GWAS and the GWAS Catalog methods for constructing the PRS involve incorporating only loci that reached genome-wide significance for at least one study, as non-significant loci are usually not reported. However, incorporating non-significant loci that are associated with the primary phenotype may help improve the predictive ability of the PRS [8, 16]. We found that incorporating additional loci that would not reach genome-wide significance did improve the PRS’ ability to discriminate cases from controls for the primary phenotype up to a point. In particular, PRS constructed using SNPs with p-values less than 5×10^−8^ or 5×10^−7^ resulted in the best performance, but further increasing the p-value threshold resulted in reduced performance. Crucially, we also observed stronger associations between the PRS and secondary phenotypes for PRS constructed using depths of 5×10^−8^ and 5×10^−7^. These results suggest that some benefit may be seen by incorporating loci that do not reach significance into the PRS construction but incorporating too many loci with larger p-values may not improve the predictive ability of the PRS (for both primary and secondary phenotypes). However, this gain or reduction in PRS performance may depend on the phenotype of interest and on the prevalence of the phenotype in the analytical sample.

As a product of this study, we provide an online visual catalog *PRSweb* that provides PRS-PheWAS results for the various skin cancer phenotypes for PRS constructed using the different methods explored in this paper. *PRSweb* will provide a routine way to compare different PRS construction methods and to explore PRS-PheWAS results in detail. Additionally, *PRSweb* provides the PRS construction details, which researchers can download and use in their own analyses. In the future, we plan to extend this online platform to include PheWAS for many other cancer phenotypes, which will make this online platform a general tool for identifying phenotypes related to particular types of cancer.

One limitation of the generalizability of this study comes from the homogeneous race profile of MGI and UK Biobank. UK Biobank consists of subjects of primarily European descent, and we restricted our analyses to subjects of European descent in MGI (excluding about 10% of the subjects in MGI) in order to ensure greater comparability between the two datasets. Additionally, many of the existing GWAS were conducted on European populations, and we wanted to consider similar samples when comparing the performance of PRS constructed using summary statistics from European populations. Unlike UK Biobank, MGI is not a population-based sample; rather, it is a sample of patients recruited from a large academic medical center. Patients were recruited prior to surgery through the anesthesiology department, and therefore they may present a potential for selection bias. Additionally, the comparative performance of the PRS across construction methods will depend on the phenotype of interest. In spite of these limitations, a principled comparison of the various methods explored in this paper may provide researchers with a sense of the robustness of their PheWAS inference to the PRS construction method and an analytical framework for exploration of shared genetic architecture of related traits.

## Acknowledgement

The authors acknowledge the University of Michigan Medical School Central Biorepository for providing biospecimen storage, management, and distribution services in support of the research reported in this publication. The presented research was funded by the National Cancer Institute at the National Institutes of Health (P30 CA046592 [LGF, LJB, MS, BM] and T32 CA83654 [RBP]). Part of this research has been conducted using the UK Biobank Resource under application number 24460. This material is based upon work supported by the National Science Foundation under Grant No. (NSF DMS 1712933). Any opinions, findings, and conclusions or recommendations expressed in this material are those of the author(s) and do not necessarily reflect the views of the National Science Foundation.

## Web resources

University of Michigan Medical School Central Biorepository; https://research.medicine.umich.edu/our-units/central-biorepository

UK Biobank; http://www.ukbiobank.ac.uk/

UK Biobank GWAS summary statistics; https://tinyurl.com/UKB-SAIGE

TOPMed variant browser, https://bravo.sph.umich.edu/freeze5/hg38/

TOPMed program, https://www.nhlbi.nih.gov/science/trans-omics-precision-medicine-topmed-program

Minimac4; https://genome.sph.umich.edu/wiki/Minimac4

BCFtools; https://samtools.github.io/bcftools/bcftools.html

KING; http://people.virginia.edu/∼wc9c/KING/

FASTINDEP; https://github.com/endrebak/fastindep

PLINK; https://www.cog-genomics.org/plink2/

Eagle; https://data.broadinstitute.org/alkesgroup/Eagle/

UCSC Genome Browser; http://genome.ucsc.edu/

R; https://cran.r-project.org/

NHGRI-EBI GWAS Catalog; https://www.ebi.ac.uk/gwas/

dbSNP; https://www.ncbi.nlm.nih.gov/projects/SNP/

Imputation server; https://imputationserver.sph.umich.edu/

Jinja, https://github.com/pallets/jinja

Locuszoom, https://github.com/statgen/locuszoom

PRSweb; https://statgen.github.io/PRSweb

## Supporting information

**S1 File. Supporting Material.** This file contains supporting Figures A-J and Tables A-E.

**S2 File. Supporting Tables.** This Excel file contains the following tables

- Sheet 1: Table F, ICD9 codes to PheWAS code translations
- Sheet 2: Table G, ICD10 codes to PheWAS code translations
- Sheet 3: Table H, MGI and UK Biobank Phenome
- Sheet 4: Table I, Risk SNP Selection
- Sheet 5: Table J, Risk SNP Selection Depth
- Sheet 6: Table K, Omnibus Table Significant Results

